# Slow but not fast theta tACS modulation of hippocampal-cortical connectivity compresses temporal distance judgments

**DOI:** 10.64898/2026.11.15.766744

**Authors:** Yuejuan Wang, Ying Jing, Laura Ivacz, Felix Duecker, Peter De Weerd, Vincent van de Ven

## Abstract

The temporal organization of episodic memory is thought to depend critically on the hippocampus, potentially coordinated by theta-band oscillations. Here, we used concurrent transcranial alternating current stimulation (tACS) and functional magnetic resonance imaging (fMRI) to examine whether slow theta stimulation (3 Hz) applied over the left parietal cortex modulates hippocampal–cortical functional connectivity during memory encoding, and how this relates to subsequent temporal memory performance. Twenty-seven participants encoded sequences of object-scene pairs that were presented either within a single context or across a context boundary, under different tACS conditions (3 Hz, 8 Hz, and no stimulation), and afterwards judged the temporal distance between item pairs. Behaviorally, 3 Hz stimulation led to significantly compressed temporal distance judgments compared to 8 Hz, suggesting altered temporal memory reconstruction. Further, 3 Hz tACS selectively reduced hippocampal connectivity with occipital cortex, comprising left calcarine and lateral occipital clusters, and reduced hippocampal activity during encoding. In contrast, 8 Hz tACS reduced connectivity between the stimulation site and right early visual cortex. Neither frequency modulated hippocampal-cortical connectivity during retrieval. These results suggest that slow theta oscillations can shape hippocampal-cortical network dynamics in a frequency-specific manner, influencing the construction of temporal relations in memory.

**Highlights:**

- Concurrent tACS–fMRI with a focal concentric-ring montage over left parietal cortex (P3) allowed frequency-specific theta stimulation (3 Hz, 8 Hz, no stimulation) to be delivered *during* encoding with per-participant SimNIBS E-field modeling confirming focal targeting
- Slow (3 Hz) but not fast theta (8 Hz) tACS compressed subsequent temporal distance judgments
- Slow but not fast theta tACS selectively reduced left-hippocampal functional connectivity with visual cortex during encoding
- 3 Hz tACS also reduced hippocampal BOLD amplitude without changing visual-cortex activity. The size of reduction correlated with individually simulated E-field strength at P3, in line with indirect, network-mediated modulation rather than direct stimulation
- Fast but not slow theta tACS reduced parietal-occipital connectivity, suggesting a frequency-based dissociation of task-related neural pathways

## Introduction

The ability to perceive and remember the temporal relationships between events is fundamental to human cognition. Temporal memory allows us to reconstruct the order and spacing of past experiences, which is critical for learning, decision-making and anticipating future outcomes (Clewett & Davachi, 2017; Eichenbaum, 2017; Tsao et al., 2022). A prominent theoretical framework for understanding temporal memory is event segmentation theory, which posits that the brain organizes continuous experiences into discrete episodes, or “events,” based on changes in perceptual and contextual information (Heusser et al., 2018; Kurby & Zacks, 2008; Reynolds et al., 2007; Swallow et al., 2009; Zacks et al., 2007). These event boundaries structure how temporal relationships are encoded and later retrieved, producing systematic biases in temporal memory judgments.

One consequence is that event boundaries affect how we perceive or remember the temporal distance between items presented in a series. Participants tend to report shorter temporal distances between items if they were presented in the same segmented context than if an event boundary occurred between them during presentation (Ezzyat & Davachi, 2014; Rouhani et al., 2020; Wang & Lapate, 2025; Wen & Egner, 2022). The hippocampus may play a role in how boundaries affect temporal context recollection. Functional magnetic resonance imaging (fMRI) studies showed that hippocampal activity patterns for individual items during encoding were more similar when the items were presented within the same context, compared to when they crossed a boundary (Ezzyat & Davachi, 2014, 2021; Hsieh et al., 2014). Further, hippocampal pattern similarity at encoding was higher for items that participants subsequently rated as temporally close than for those rated as further apart, indicating that hippocampal pattern similarity predicted temporal proximity judgments (Ezzyat & Davachi, 2014). These findings fit with a growing consensus that hippocampal function supports encoding and representation of temporal contexts (Eichenbaum, 2017; Ranganath & Hsieh, 2016; Tsao et al., 2022).

Importantly, theta oscillations (3-8 Hz), a hallmark of hippocampal activity, have been implicated in coordinating memory encoding and event segmentation (Griffiths & Fuentemilla, 2020; Gupta et al., 2012; Heusser et al., 2016). For example, Gupta et al. (2012) provided evidence that theta-gamma coupling in rodent hippocampus may reflect how spatial boundaries in a maze trajectory are temporally represented in the brain. They found that gamma activity phase-locked to theta oscillations tended to predict future locations when an animal was entering a new spatial segment and to reflect previous locations when approaching the end of a segment. This suggests a dynamic encoding of spatial context aligned with contextual boundaries. Notably, their analyses also indicated that spatial segments defined by boundaries were associated with slower theta frequencies and a higher number of gamma bursts, implying that lower-frequency theta rhythms may support the integration of more items across extended events and may therefore enhance segmentation and temporal memory. This finding fits with a growing body of evidence suggesting that different theta frequencies may subserve distinct cognitive functions. Intracranial recordings have shown that slow theta (∼3 Hz) power in the hippocampus is selectively associated with memory encoding (Lega et al., 2012), whereas fast theta (∼8 Hz) is more closely linked to spatial navigation and non-mnemonic processes (Goyal et al., 2020). Further evidence for a slow-fast theta dissociation comes from a human intracranial EEG study showing that hippocampal-cortical slow theta synchrony was related to faster working memory responses while fast theta synchrony was related to impaired performance (Gray et al., 2026).

One promising approach to probing the causal role of theta oscillations is transcranial alternating current stimulation (tACS), a noninvasive technique capable of modulating brain oscillations in a frequency-dependent manner (van der Plas & Hanslmayr, 2020; Wischnewski et al., 2023). Several studies have shown that tACS delivered at a theta frequency can modulate associative recognition (Lang et al., 2019; Meng et al., 2021; Vulić et al., 2021; Živanović et al., 2022) while not affecting perceptual recognition (Meng et al., 2021), suggesting that theta stimulation may be functionally specific to associative memory processes. Recent tACS studies have further supported the slow-fast theta dissociation in memory, with slow theta tACS over fronto-parietal cortex enhancing working memory performance, compared to fast theta stimulation (Aktürk et al., 2022; Wang et al., 2024; Wolinski et al., 2018). We recently showed that slow (3 Hz), but not fast (8 Hz) theta tACS over parietal cortex during the presentation of lists of visual items enhanced subsequent temporal memory accuracy for those items (Wang et al., 2024). These findings indicate that frequency-specific electrical stimulation can be used to dissociate different temporal mnemonic functions within the theta range. Moreover, there is growing evidence that tACS delivered over parietal cortex can modulate hippocampal activity or hippocampal-cortical connectivity in the monkey (Krause et al., 2019) and human brain non-invasively (Kaiser et al., 2025; Luo et al., 2025), possibly by utilizing anatomical (Niu & Palomero-Gallagher, 2023; Vieira et al., 2025) or functional connectivity between parietal cortex and hippocampal areas (Vincent et al., 2006; Wang et al., 2014).

In the present study, we investigated if slow (3 Hz) but not fast (8 Hz) theta tACS influences temporal memory and event segmentation by modulating hippocampal network activity during simultaneously measured fMRI. We administered tACS over parietal cortex based on our previous findings of modulating associative memory (Meng et al., 2021; Wang et al., 2024) and modulating hippocampal-cortical connectivity (Kaiser et al., 2025). Given the hippocampus’s central role in representing temporal context and detecting event boundaries, we hypothesized that 3 Hz, but not 8 Hz, tACS modulates hippocampal function and hippocampal-cortical connectivity relevant to the task. Further, we hypothesized that 3 Hz, but not 8 Hz, tACS alters temporal memory performance.

## Methods

### Participants

Twenty-seven healthy participants (17 females, mean (SD) age=26.74 (4.78)) were recruited from Maastricht University. An a priori power calculation in G*Power using effect sizes from previous temporal distance studies (Ezzyat & Davachi, 2014; McClay et al., 2023; alpha=0.05, effect size=0.8, power=0.9, two-tailed) suggested a minimum sample size of N=19. We included more participants to obtain better statistical power for comparison between brain stimulation conditions. Before visiting the lab, participants completed screening forms to assess their eligibility for transcranial electric stimulation and magnetic resonance imaging (MRI). On the day of the experiment, participants reviewed and signed the screening forms again on-site and provided informed consent before the study commenced. The study was approved by the Ethical Review Committee of Psychology and Neuroscience (ERCPN) at the Faculty of Psychology and Neuroscience, Maastricht University. Participants received financial compensation of 20 EUR for their participation.

### Procedures

Before entering the scanner, participants practiced the behavioral tasks outside of the MRI environment. They then completed another practice session inside the scanner during the anatomical measurements. The experiment consisted of three functional runs, each including three repetitions of the encoding and memory tasks (Figure 1). The order of the three tACS conditions was presented pseudo-randomly to each participant, such that each condition was presented once in a functional run and orders did not repeat across runs for a participant. Thus, each tACS condition was repeated three times for each participant throughout scanning. Tasks and stimulation waveforms were programmed in PsychoPy 3 (Peirce et al., 2019).

**Figure 1.**
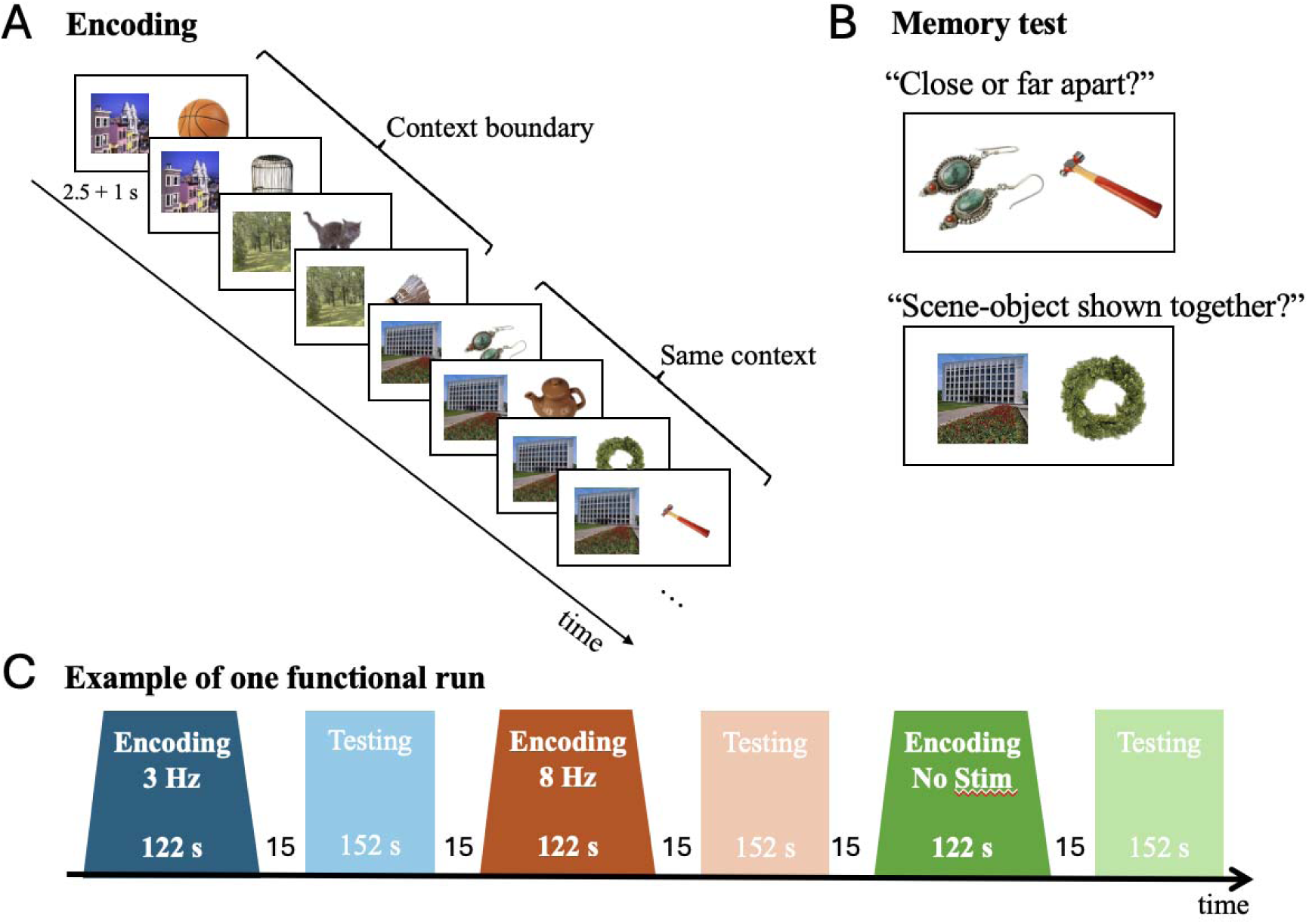
Experimental design. The behavioral tasks included an associative encoding task (A) and memory testing tasks (B) for temporal distance and scene-object associations. C: In each of three functional MR runs, there were three blocks of encoding and testing tasks. Three tACS conditions (no stimulation, 3 Hz and 8 Hz) were randomly assigned among the three encoding phases. Numbers indicate seconds duration for the various task blocks or resting periods between blocks.

During each encoding task, participants were presented with eight quartets of object-scene pairs, totaling 32 item-scene pairings. Within each quartet, scene presentation varied depending on the condition: in half of the quartets, all four objects were paired with the same scene (same-context condition), while in the other half, the scene changed after the second object (boundary condition). Each object-scene pair was displayed for 2.5 seconds, with the scene positioned on the left and the object on the right (Figure 1A). Participants were instructed to imagine encountering the object within the given scene and rate the ease of this mental simulation on a scale from 1 (very easy) to 4 (very difficult). The rating scale was displayed alongside the item-scene pair and disappeared as soon as participants made their response. They had up to 3.5 seconds to respond before the next pair appeared. The order of scene and object presentations was randomized across participants.

After each encoding task, participants completed a temporal distance judgment task (Figure 1B). In this task, test pairs were selected from the object images shown during encoding. Each test pair consisted of objects that had appeared either in the first and last positions of a quartet (long distance) or in the second and third positions (short distance). Participants were asked to recall the perceived temporal distance between the two items (“how far apart in time do you think the items were presented during encoding”) and rate it on a four-point scale from 1 (very close) to 4 (very far) within a 4.7-second response window. The rating scale disappeared upon button press and was followed by a brief 0.3-second blank screen before participants proceeded to the next trial. The test pairs were presented in a randomized order across participants, with each session including a total of 16 trials.

After completing the temporal distance task, participants performed an associative memory task (Figure 1C). In this task, they were presented with an object-scene pair and asked to determine whether the object had originally been paired with that scene during encoding. For each test trial, the third item from a quartet was shown either with the same scene it had been paired with during encoding (matching pair) or with a scene from a different quartet (non-matching pair). Participants responded within a 2.7-second window using a four-point scale: 1 (Sure Yes), 2 (Maybe Yes), 3 (Maybe No), and 4 (Sure No). Each trial was separated by a 0.3-second intertrial interval. Each session included a total of eight trials, with an equal number of match and non-match trials (four each). The order of test trials was randomized across participants.

Each participant was presented with nine learning lists each consisting of 32 objects and 12 scenes. The experiment included three functional runs, with each run containing three encoding and testing cycles. Items and scenes were not repeated within a participant.

### Behavioral data analysis

The data were processed in Matlab (https://www.mathworks.com) with customized code and analyzed statistically in jamovi (https://www.jamovi.org). For encoding response time, a repeated measures ANOVA was used with tACS condition (no stimulation, 3 Hz and 8 Hz), context (same context vs. context boundary) and encoding position (1,2,3,4) within each quartet as within-subject factors. For the temporal distance task, we used a repeated measures ANOVA to analyze each of two dependent variables, response time and the distance rating, with tACS condition, context and distance (short vs. long) as within-subject factors. For the associative memory task, we calculated d’ as a signal detection theory measure of memory sensitivity from hit rates and false alarm rates (Stanislaw & Todorov, 1999). Hit and false alarm rates of 0 or 1 were corrected respectively by adding 0.5 or −0.5 to the numerator when calculating the proportions. Statistical results were deemed significant if the ensuing p-value was below the alpha threshold of 0.05.

### Transcranial alternating current stimulation (tACS)

Before scanning, tACS electrodes were positioned on the participant’s scalp (Figure 2A). The stimulation was delivered by a NeuroConn DC STIMULATOR TES device at the P3 location, identified using the 10-20 electrode system with an EEG cap. The electrode placement was marked on the scalp with an eyebrow pencil to ensure consistency. A ring-shaped electrode (the inner circular electrode: 2 cm diameter; the outer electrode: 10 cm diameter; width 1.2 cm) was used to provide focal stimulation (Bortoletto et al., 2016; Kaiser et al., 2025). To minimize impedance, the scalp was first cleaned with alcohol, after which a conductive paste was applied to the electrode and skin. Care was taken to ensure that the paste of the two electrodes did not come into contact. Impedance levels were lowered to below 5 kΩ for all participants, after which participants underwent a brief familiarization session with tACS stimulation.

**Figure 2.**
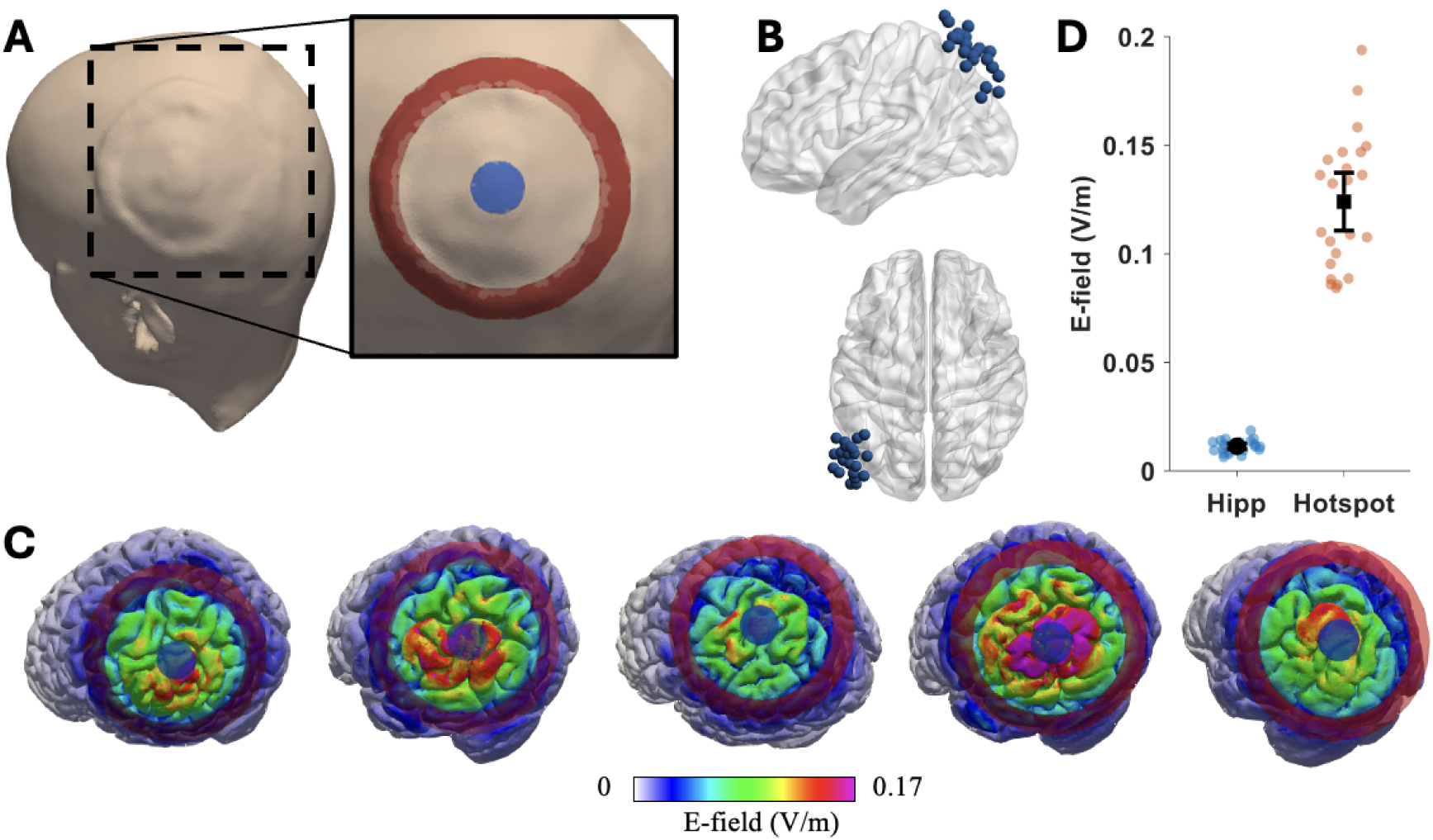
Concentric-ring tACS electrode layout and simulated electric field distribution. A: An example of an individual head model reconstructed from the structural MRI, with the contour of the concentric-ring electrodes modelled in SimNIBS superimposed on the head model (inset). B: Distribution of stimulation targets (centers of the central electrodes) in MNI space across participants. C: Simulated electric-field (E-field) magnitude distribution on the cortical surface viewed from a lateral perspective to illustrate the spatial pattern of the induced field in five representative participants (leftmost representation is the same participant as in A). All maps are shown in the same color range (E-field magnitude in V/m) to allow direct comparison across participants. D: Distribution of simulated E-fields for left hippocampus (Hipp) and central electrode hotspots. Error bars indicate 95% confidence intervals.

During each functional run, the order of the three tACS stimulation conditions was randomized (Figure 1D). Stimulation during the 3 and 8 Hz conditions was delivered at a constant current strength of 2 mA (peak-to-peak) for a period lasting 112 seconds, which was flanked by a gradual ramp-up and ramp-down of 5 seconds at the beginning and end of each condition. No electrical stimulation was delivered during the no stimulation condition. Between stimulation conditions, there was a 182 second interval, which included a 15 second break before and after the testing tasks.

For concurrent tACS-fMRI, we implemented the same tACS setup as described in Kaiser et al. (2025), with the stimulator placed in the console room and an MR-compatible system delivering stimulation inside the scanner. The stimulation waveform protocol was designed in PsychoPy/Python, specifying the precise timing of ramping phases, stimulation duration and frequency for each condition. This waveform protocol was then transmitted to the output device via a digital-to-analog converter (National Instruments, Austin, Texas, USA).

### Electric field modeling

For each participant, electric field (E-field) simulations were performed on the individual head mesh using the finite element method implemented in SimNIBS v4.1.0 (Saturnino et al., 2019; Thielscher et al., 2015). Individual head model reconstruction was performed using the CHARM pipeline, which integrates FreeSurfer outputs to improve the representation of fine anatomical structures, particularly small sulci (Puonti et al., 2020). The resulting finite element head models consisted of approximately 7.3 × 10 nodes and 4.1 × 10 tetrahedral elements, with an average element volume of approximately 0.57 mm³ in the cortical regions. Ten tissue types were included in the models with standard isotropic conductivity values: white matter (□WM=0.126 S/m), gray matter (□GM=0.275 S/m), cerebrospinal fluid (□CSF=1.654 S/m), bone (□B=0.01 S/m), scalp (S=0.465 S/m), eyeballs (□EB=0.500 S/m), compact bone (□CB=0.008 S/m), spongy bone (SB=0.025 S/m), blood (□BL=0.600 S/m) and muscle (□M=0.160 S/m). All tissues were modeled as isotropic.

E-field simulations were conducted using a concentric-ring electric stimulation montage with the central electrode (diameter=20 mm) and surround electrode (outer diameter=100 mm, inner diameter=76 mm) positioned over left P3. Electrode thickness was set to 2 mm and electrode conductivity set to 8 S/m. A current intensity of ±1 mA was applied (−1 mA at the central electrode, +1 mA at the ring electrode). For each participant, the resulting E-field magnitude distribution was computed across the whole brain.

To quantify stimulation intensity at the cortical target, the peak E-field magnitude location was identified for each participant (typically located underneath the central electrode). A spherical region of interest (ROI, diameter=8 mm) centered on the peak location was defined and constrained to gray matter. Mean E-field magnitude within this ROI was extracted for subsequent analyses. Additionally, region-wise E-field values were extracted using the Automated Anatomical Labeling (AAL) atlas, allowing correlation analyses between E-field strength and behavioral or neuroimaging measures.

### MRI acquisition

Anatomical and functional imaging were performed using a 3T Magnetom Prisma Fit scanner (Siemens, Erlangen, Germany). The session began with a fast T2-HASTE scan (9 slices) to localize the hippocampus. A high-resolution T1-weighted MP-RAGE sequence was then used to obtain anatomical reference images (TR/TE=2250/2.21 ms, voxel size=1 mm³ isotropic, FOV=256 × 256 mm, 192 slices). The anatomical scan lasted approximately 5 minutes, followed by three functional runs. Each functional run lasted 15.5 minutes, during which functional data were acquired using a T2-weighted echo-planar imaging (EPI) sequence (TR/TE=1000/30 ms, voxel size=2.6 × 2.6 × 3 mm, FOV=159 × 129 mm, A-P × R-L, 42 slices, 930 volumes, multi-band factor=3).

### MR image pre-processing

Functional and anatomical data were preprocessed using a modular preprocessing pipeline implemented in the CONN toolbox (v.21.a), which incorporates image processing routines from SPM12. The pipeline included realignment with correction for susceptibility distortion interactions, slice timing correction, outlier detection, segmentation, normalization to MNI space, and spatial smoothing.

First, functional images were realigned using SPM’s realign & unwarp procedure, in which all scans were coregistered to the first volume of the first functional run using a six-parameter (rigid body) transformation. This step also corrected for motion-related susceptibility distortions using B-spline interpolation. Next, slice timing correction was performed using sinc temporal interpolation to align all slices to a common mid-acquisition time, accounting for the interleaved acquisition order. Outlier scans were identified using the Artifact Detection Toolbox (ART) and were excluded based on framewise displacement (>0.9 mm) or global signal changes exceeding five standard deviations.

Following motion correction, functional and anatomical images were segmented into gray matter, white matter, and cerebrospinal fluid (CSF) and were normalized to MNI space using SPM’s unified segmentation and normalization algorithm, with tissue probability maps from the IXI-549 template. The normalized functional images were resampled to 2 mm isotropic voxels and smoothed using a Gaussian kernel of 4 mm full width at half maximum (FWHM).

To reduce physiological and motion-related noise, functional data underwent a standard denoising procedure. This included the regression of five CompCor noise components from white matter and CSF, motion parameters and their first-order derivatives (12 factors), outlier scans (up to 75 factors), session and task effects with their first-order derivatives (18 factors), and linear trends (2 factors) within each functional run. A bandpass filter (0.008–0.09 Hz) was applied to the BOLD signal to further minimize high-frequency noise.

### Functional connectivity analyses

The impact of tACS on functional connectivity was assessed using seed-based correlation (SBC) analysis, which measures the timepoint-by-timepoint Pearson correlation between the preprocessed BOLD time series of a designated ROI and the time series of all other voxels in the brain. The left hippocampus, identified from the Harvard-Oxford brain atlas, served as the seed region. Prior to analysis, functional runs were normalized and concatenated to create a continuous time series for each participant. The statistical approach involved two levels of analysis. At the first level, individual correlation maps were generated for each seed and participant, separately for three tACS conditions and the encoding and temporal memory retrieval tasks, yielding six connectivity maps per participant corresponding to different tACS conditions during encoding and retrieval. Each correlation map was then normalized using Fisher’s r-to-z transformation.

At the second level, statistical contrasts were defined at the subject level to examine differences in connectivity patterns across conditions. Following our hypothesis, we tested if 3 Hz stimulation affected hippocampal-seeded connectivity relative to the no stimulation and 8 Hz condition, using a within-subject contrast *c*=[0.5, −1, 0.5] for the no stimulation, 3 Hz and 8 Hz conditions. We also tested if 8 Hz stimulation affected connectivity using the contrast *c*=[0.5, 0.5, −1]. The t-test statistical maps were initially thresholded (uncorrected *p*-value of 0.001 and cluster level FDR-correction, *q*=0.05) prior to visualization on a 3D surface rendering of the MNI template. Post-hoc pairwise comparisons of correlation values across tACS conditions were subsequently conducted in significant voxel clusters to describe the pattern of results (i.e., treating significant voxel clusters as omnibus test results).

To assess “baseline” hippocampal-cortical connectivity during task performance (i.e., without tACS), we conducted a left hippocampus seed-based analysis in no stimulation condition during encoding (*c*=[1, 0, 0]). We also assessed baseline hippocampal-cortical connectivity during temporal memory testing using the same statistical contrast.

### Hippocampal activity

In left hippocampus, denoised voxel-wise BOLD signals were averaged to obtain an ROI-level time series. Based on condition onsets and durations, time points corresponding to each experimental condition (no stimulation, 3 Hz, 8 Hz) were extracted from the full time series. We then calculated percent signal change for each condition by averaging signal amplitude and dividing by a baseline of the first 5 seconds of each condition, and averaged the percent signal change amplitudes across the three runs of each condition.

## Results

### Simulated electric field

Four participants were excluded because of poor task performance, reported poor vision during scanning, or failure of the experimental setup to log task responses during scanning, leaving N=23 participants for the final analysis. Figure 2 illustrates the concentric-ring tACS electrode configuration and the corresponding simulated electric-field distributions. The E-field simulations showed that the induced electric field was strongest directly beneath the central electrode positioned over the left parietal cortex (around the P3 location), with the field intensity gradually decreasing toward surrounding cortical regions (Figure 2B). This spatial pattern indicates that the concentric-ring montage produced a relatively focal stimulation profile. The distribution of stimulation targets in MNI space across participants further confirmed that electrode placement was spatially consistent and clustered around the left parietal cortex (Figure 2C). Figure 2D shows distributions of average E-field values for the left hippocampal ROI (mean V/m=0.011, 95% *CI* = [0.010, 0.012]) and the P3 hotspots (mean=0.124, 95% *CI* = [0.113, 0.135]).

### Encoding response time

Figure 3A illustrates the encoding response time as a function of encoding position and context. The three-way repeated measures ANOVA (position, context, tACS) revealed a significant main effect of position (*F*(3, 66)=51.90, *p*<0.001, η*_p_^2^*=0.70) and of context (*F*(1, 22)=12.88, *p*=0.002, η*_p_^2^*=0.37) and a significant position × context interaction effect (*F*(3, 66)=11.96, *p*<0.001, η*_p_^2^*=0.35). In the same context condition, participants responded slower to the 1^st^ item of a quartet (mean [SE]=1.72 [0.05]) than to items at the other three positions (2^nd^=1.54 [0.05], 3^rd^=1.55 [0.04], 4^th^=1.55 [0.04]; 1^st^ vs. 2^nd^: *t*(22)=7.30, *p*<0.001, Cohen’s *d*=0.53; 1^st^ vs. 3^rd^: *t*(22)=8.92, *p*<0.001, Cohen’s *d*=0.55; 1^st^ vs. 4^th^: *t*(22)=7.83, *p*<0.001,

**Figure 3.**
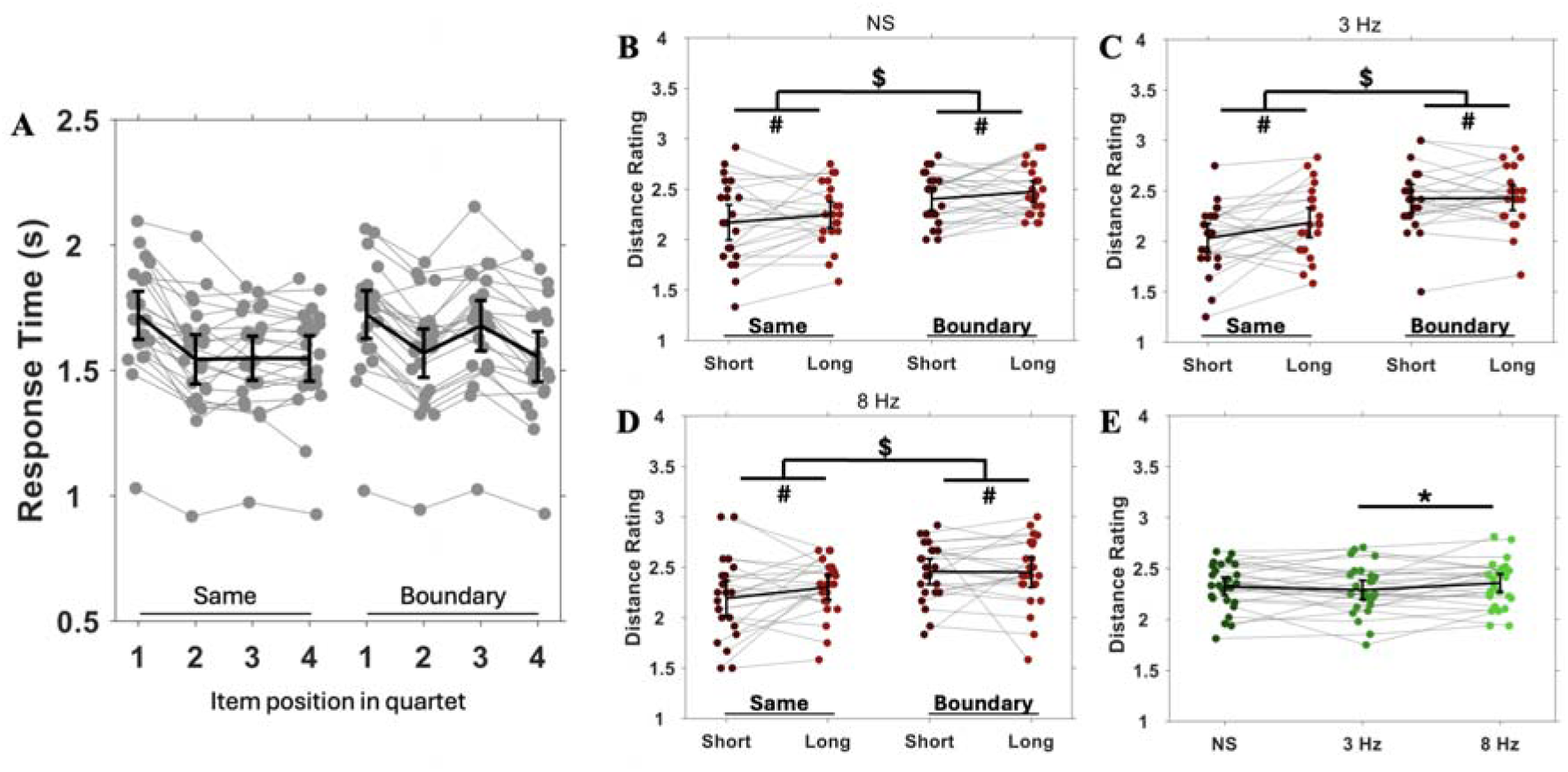
Behavioral results of encoding and temporal distance. A: Encoding response times (RT) as a function of context (same and boundary conditions) and position. B-E: Temporal distance judgments as a function of context and distance in no-stimulation (B), 3 Hz (C) and 8 Hz tACS condition (D). E: Temporal distance ratings pooled across distance and boundary conditions under each of the three tACS conditions. Error bars represent the 95% confidence interval. ^$^ main effect of the context (p<0.001), ^#^ main effect of distance (p<0.05), * p<0.05.

Cohen’s *d*=0.55). In the context boundary condition, participants responded slower to items at the 1^st^ position of a quartet (1.72 [0.05]) than the 2^nd^ (1.57 [0.05]), 3^rd^ (1.68 [0.05]) and 4^th^ position (1.55 [0.05], 1^st^ vs. 2^nd^: *t*(22)=9.67, *p*<0.001, Cohen’s *d*=0.44; 1^st^ vs. 3^rd^: *t*(22)=2.48, *p*=0.021, Cohen’s *d*=0.12; 1^st^ vs. 4^th^: *t*(22)=7.26, *p*<0.001, Cohen’s *d*=0.50), while response times for the 3^rd^ position (1.68 [0.05]) were slower than for the 2^nd^ and 4^th^ position (3^rd^ vs. 2^nd^: *t*(22)=6.34, *p*<0.001, Cohen’s *d*=0.32; 3rd vs. 4th: t(22)=6.76, *p*<0.001, Cohen’s *d*=0.38). In addition, participants responded slower (*t*(22)=7.39, *p*<0.001, Cohen’s *d*=0.42) to items at which the scene changed in the context boundary condition (i.e., 3^rd^ position, 1.68 [0.05]), compared to the 3^rd^ position in the same context condition (1.55 [0.04]), indicating slower processing when a context change occurred. The main effect of tACS was not significant (*p*=0.433), suggesting that tACS did not influence how participants responded to the boundary during encoding.

### Temporal distance task

Figure 3B-D illustrates the mean temporal distance ratings as a function of context and physical distance in the no-stimulation (Figure 3B), 3 Hz (Figure 3C) and 8 Hz tACS condition (Figure 3D). The repeated measures ANOVA revealed significant main effects of tACS (*F*(2, 44)=3.42, *p*=0.042, η*_p_^2^*=0.14), context (*F*(1, 22)=31.15, *p*<0.001, η*_p_^2^*=0.57) and distance (*F*(1, 22)=5.17, *p*=0.033, η*_p_^2^*=0.19). None of the interaction terms were significant (*p*s>0.086). The temporal distance rating (Figure 3E) was significantly lower (*t*(22)=2.34, *p=*0.029, *Cohen’s d*=0.21) during 3 Hz tACS stimulation (2.27 [0.05]) compared to 8 Hz stimulation (2.35 [0.05]). The temporal distance rating during no stimulation did not significantly differ from 3 Hz or 8 Hz. In addition, participants made significantly higher distance ratings for the pairs in the context boundary condition (2.44 [0.04]) than the pairs in the same context condition (2.19 [0.06]), which aligns with previously reported temporal compression effects for within-event trials (Ezzyat & Davachi, 2014; McClay et al., 2023; Wen & Egner, 2022). The temporal distance rating was higher for the long distance pairs (2.35 [0.04]) than the short distance pairs (2.28 [0.05]), suggesting that the true encoding distance influenced the temporal distance judgments.

A repeated measures ANOVA of response times showed no significant effects (context: *p*=0.088, distance: *p*=0.120, tACS *p*=0.514; context × distance: *p*=0.748; context × tACS: *p*=0.365; distance × tACS: *p*=0.338; context × distance × tACS: *p*=0.074).

To assess if individual differences in 3 Hz stimulation were related to temporal ratings, we correlated peak simulated E-fields under the central electrode to temporal ratings during the 3 Hz condition. The correlation was not significant (r=0.04, p=0.85). We further considered the simulated E-fields being a difference measure of stimulation against a zero-field baseline, which we correlated with the difference between 3 Hz and the no stimulation condition, and found no significant result (r=-0.0, p=0.998). Individual differences in E-fields thus did not explain subject-level variation in temporal distance performances.

### Associative memory task

Repeated measures analysis of memory sensitivity (*d’*, Figure 4A) revealed a significant main effect of context (*F*(1, 22)=10.05, *p*=0.004, η*_p_^2^*=0.31), with better performance for items in the same context condition (1.77 [0.23]) than that in the context boundary condition (1.45 [0.23]). We found a marginal main effect of tACS (*F*(2,44)=3.20, *p*=0.051, η*_p_^2^*=0.13). Post-hoc paired comparisons between tACS conditions showed that 3 Hz tACS decreased memory sensitivity compared to no stimulation (*p*=0.024) but that the differences between 3 Hz and 8 Hz (*p*=0.222) or between 8 Hz and NS (*p*=0.199) were not significant. The context × tACS interaction effect was not significant (*p*=0.928).

**Figure 4.**
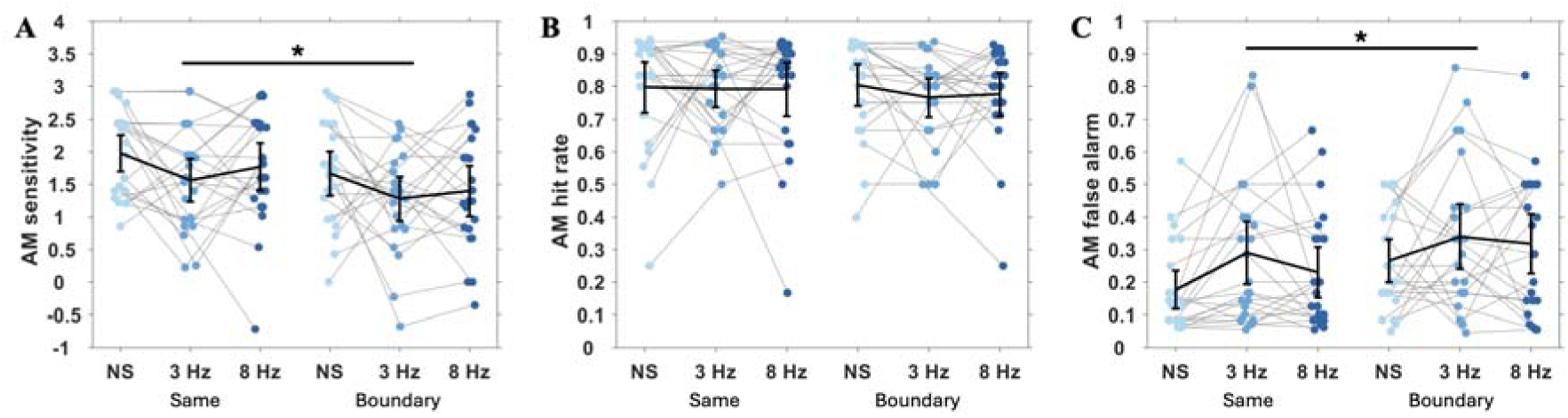
Associative memory task results. Memory sensitivity *d’* (A), hit rate (B) and false alarm rate (C) in the associative memory task as a function of context and tACS condition. Error bars represent the 95 % confidence interval.

To further examine the differences in memory performance across tACS conditions, we also analyzed hit rates and false alarm rates. The ANOVA on hit rates (Figure 4B) revealed no significant effects of tACS or context (*ps*>0.065). For the false alarm rates (Figure 4C), the repeated measures ANOVA revealed a main effect of tACS (*F*(2, 44)=4.88, *p*=0.012, η*_p_^2^*=0.18) and of context (*F*(1, 22)=6.86, *p*=0.016, η*_p_^2^*=0.24), but no significant interaction effect (*p*=0.745). The false alarm rate in the context boundary condition (0.31 [0.03]) was higher than that in the same context condition (0.23 [0.03]). In addition, average false alarm rate was significantly higher (*t*(22)=3.16, *p*=0.005, Cohen’s *d*=0.42) in the 3 Hz condition (0.32 [0.04]) than the no stimulation condition (0.22 [0.03]). The difference between 8 Hz and no stimulation condition or the difference between 3 Hz and 8 Hz (0.27 [0.04]) tACS conditions was not significant. These results suggest that associative memory performance was influenced by contextual continuity, with better memory sensitivity observed for items within the same context compared to those across a boundary, and that tACS modulated associative memory sensitivity through false alarm rates. Specifically, participants exhibited higher false alarm rates in the context boundary condition than in the same context condition, indicating that event boundaries may introduce greater memory interference. Additionally, 3 Hz tACS led to a significant increase in false alarms compared to the no-stimulation condition, suggesting that slow theta stimulation may influence associative memory by altering the specificity of retrieved information.

The repeated measures ANOVA for response times revealed no significant effects (context: *p*=0.293; tACS: *p*=0.293; context × tACS: p=0.633).

### Hippocampal-cortical connectivity during no-stimulation encoding

The hippocampal-cortical connectivity during encoding in the no stimulation condition revealed a widespread brain network (Figure 5A), which included bilateral medial and lateral parietal cortex, anterior temporal poles, medial temporal lobe areas including hippocampus and amygdala and ventromedial prefrontal cortex. A similar widespread network was found for the no stimulation condition during the memory task (Figure 5B). This network aligns with previously reported networks of cortical and subcortical regions underlying memory formation (Eichenbaum, 2017; Ranganath & Ritchey, 2012; Vincent et al., 2006).

**Figure 5.**
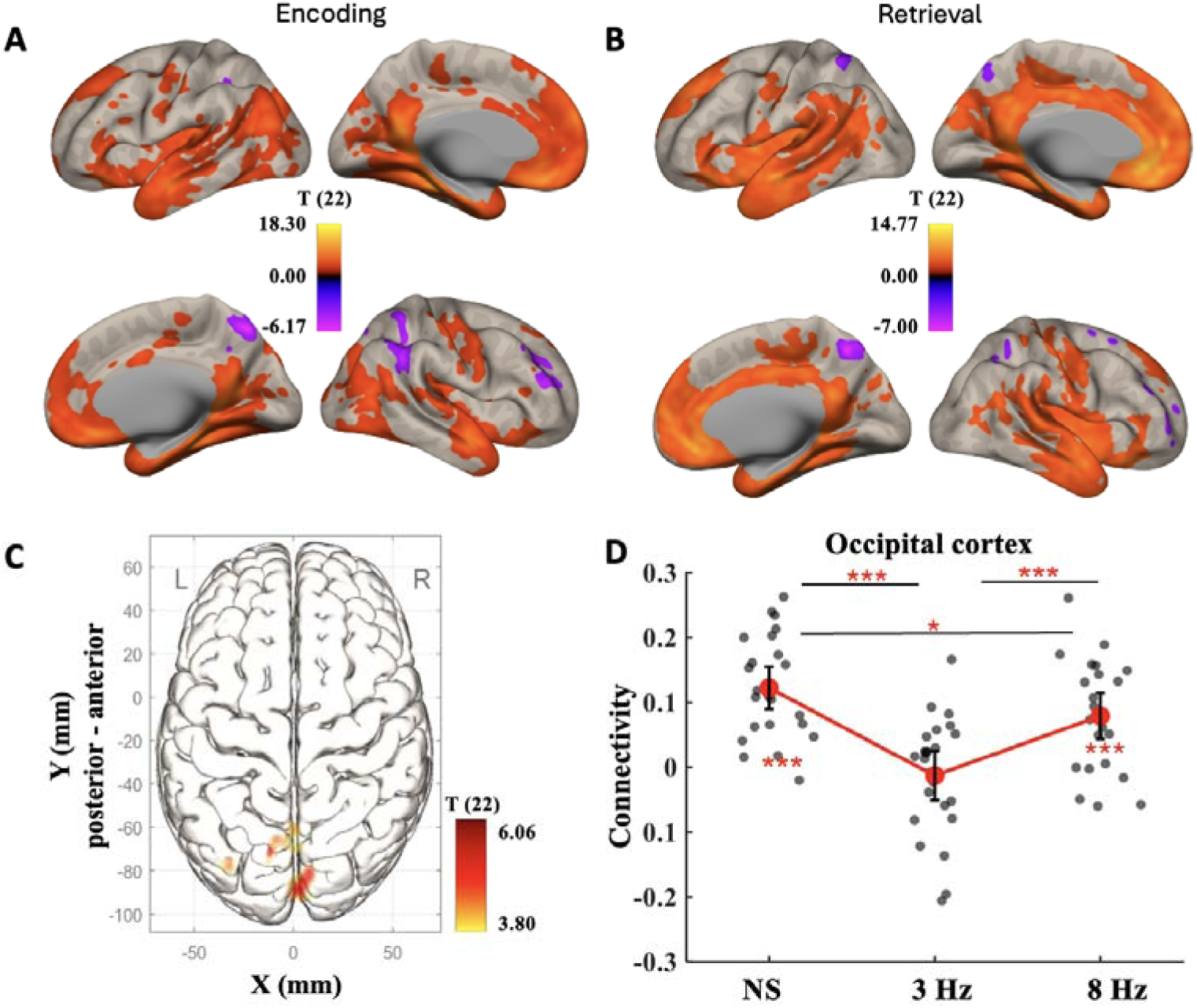
Hippocampal connectivity results. Hippocampal-cortical connectivity during the no stimulation condition in the encoding (A) and memory retrieval task (B). C: During encoding, 3 Hz tACS decreased hippocampal connectivity with left visual cortex, compared to no stimulation and 8 Hz conditions. D: Distribution of connectivity values for the three tACS conditions from the clusters shown in C. Error bars represent 95% confidence intervals. NS, no stimulation condition; * p<0.05; *** p<0.001.

### The effect of tACS on hippocampal connectivity during encoding

The hippocampal seed-based connectivity analysis for the effect of 3 Hz stimulation compared to 8 Hz and NS revealed three significant clusters in right calcarine cortex (size=116 mm³, MNI coordinates x,y,z=+2, −88, +10), left calcarine and precuneus (size=71 mm³, MNI coordinates x,y,z=-12, −72, +12) and left lateral occipital cortex (size=28 mm³, MNI coordinates x,y,z=-32, −78, −16). Figure 5C depicts the three visual cortical clusters and Figure 5D shows the distribution of connectivity values for the three tACS conditions (averaged across the three visual cortical clusters). The hippocampal-visual cortex connectivity values in the no stimulation and 8 Hz stimulation conditions were higher than 0 (no stimulation=0.12 [0.02]: t(22)=7.36, *p*<0.001, Cohen’s *d*=1.53; 8 Hz=0.08 [0.02]: t(22)=4.38, *p*<0.001, Cohen’s *d*=0.91) but decreased significantly during 3 Hz stimulation (−0.01 [0.02]; 3 Hz – no stimulation: t(22)=-8.35, 95% *CI*=[−0.17, −0.10], *p*<0.001; 3 Hz – 8 Hz: t(22)=-5.85, 95% *CI*=[−0.12, −0.06], *p*<0.001). Further, 8 Hz stimulation decreased the connectivity compared to no stimulation (t(22)=-2.27, 95% *CI*=[−0.08, −0.01], *p*=0.033). These results suggest that 3 Hz tACS significantly disrupts hippocampal connectivity with the occipital cortex, a region involved in visual processing. When analyzing seed-based connectivity for the effect of 8 Hz stimulation, we did not find any significant clusters.

### The effect of tACS on percent signal change during encoding

We next examined percent signal change activity in left hippocampus during the encoding phase. Compared to no stimulation, 3 Hz tACS significantly reduced hippocampal activity (*t*(22)=−4.10, 95% *CI* [−0.37, −0.13], *p*<0.001). In contrast, 8 Hz tACS showed no significant difference from the no stimulation condition (*p*=0.36). To directly compare frequency-specific effects, we tested the interaction contrast ((3 Hz – NS) − (8 Hz – NS)), which was significant (*t*(22)=−2.10, 95% *CI* [−0.86, −0.005], *p*=0.047), indicating that 3 Hz tACS induced a stronger modulation of hippocampal activity than 8 Hz tACS (Figure 6A). When examining activity in the visual cortex clusters (Figure 6B), we found no significant effects for either the main comparisons (3 Hz or 8 Hz vs. no stimulation) or the interaction contrast (*p*s>0.11).

**Figure 6.**
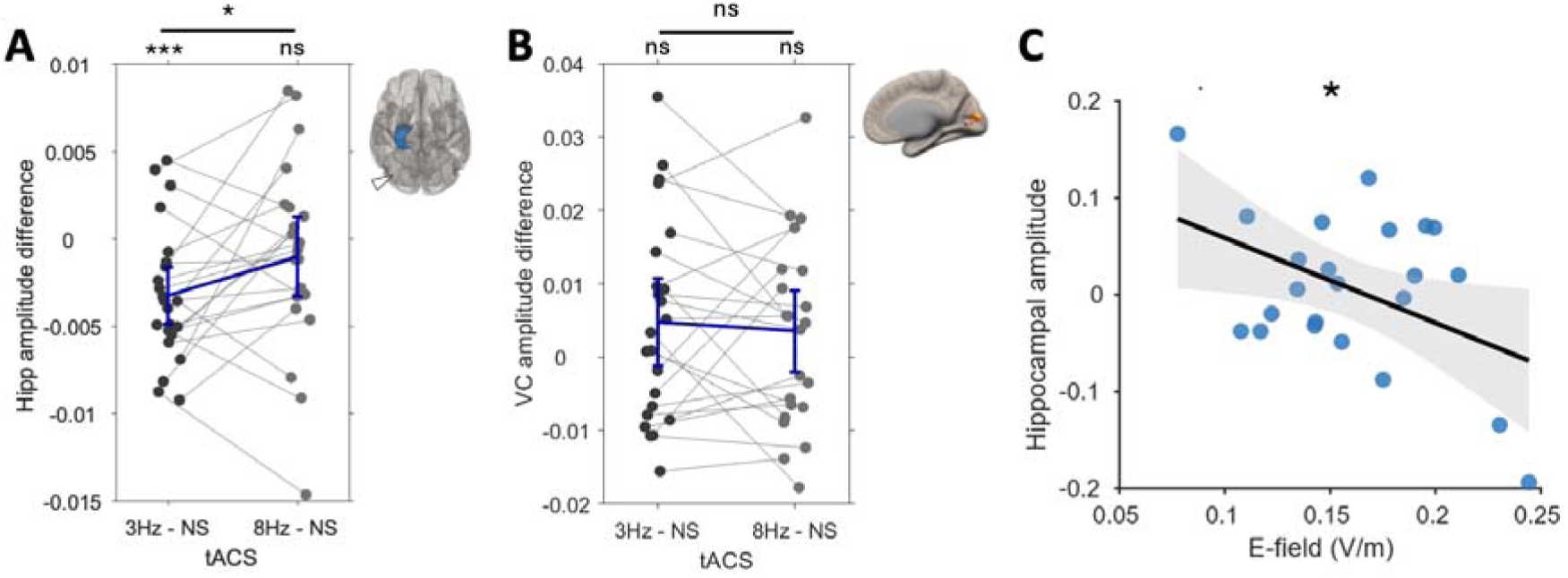
tACS-induced activity change. Amplitude change between 3 Hz or 8 Hz and no stimulation in left hippocampus (A) and visual cortex (B). Error bars represent the 95% confidence interval. C: Scatter plots showing the association between simulated E-field strength at the stimulation target and left hippocampal amplitude across participants under 3 Hz tACS. Shaded area indicates 95% confidence interval. NS, no stimulation condition; ns, not significant; * p<0.05; *** p<0.001.

We then tested if the individual variability in hippocampal response under 3 Hz tACS was related to the strength of the induced electric field, as simulated with SimNIBS (Figure 6C). We found a significant negative correlation between E-field strength at inferior parietal cortex and hippocampal amplitude (r=−0.45, p=0.031). We note here that this result does not imply that 3 Hz tACS affected hippocampal function directly, as the induced electric field is too weak. Rather, we suggest that a stronger E-field over parietal cortex is better able to modulate hippocampal function indirectly via intrinsic connectivity dynamics.

### P3-seeded connectivity during encoding

To explore brain connectivity changes in relation to the P3 location of the central electrode, we created a regional sphere of 8 mm radius at the central electrode location as seed region for a seed-based connectivity analysis, using the same statistical thresholding as for the hippocampal seed analysis. For 3 Hz stimulation, the analysis revealed no significant clusters. For 8 Hz stimulation, we found a significant cluster (size=120 mm³, MNI coordinates: x=+16, y=-72, z=+10) located in the right intracalcarine cortex (Figure 7). Post-hoc one-sample t-tests showed that the connectivity values during the no stimulation (0.001 [0.018], *t*(22)=0.07, *p*=0.943) and 3 Hz stimulation (−0.006 [0.024], *t*(22)=-0.27, *p*=0.789) were not significantly different from 0. However, connectivity during 8 Hz stimulation was significantly below 0 (−0.127 [0.019], *t*(22)=-6.48, *p*<0.001, Cohen’s *d*=-1.35) and differed significantly from connectivity during no stimulation (*t*(22)=-6.41, 95% *CI*=[−0.17, −0.09], *p*<0.001) and 3 Hz stimulation (*t*(22)=-5.31, 95% *CI*=[−0.17, −0.07], *p*<0.001). These findings suggest that 8 Hz tACS selectively decreased P3 connectivity with the right intracalcarine cortex. Notably, the absence of a 3 Hz effect on parietal-visual cortical connectivity, while 3 Hz did modulate hippocampal-cortical connectivity, points to a partial dissociation of slow and fast theta frequencies respectively targeting different neural pathways, when delivered over the same cortical location.

**Figure 7.**
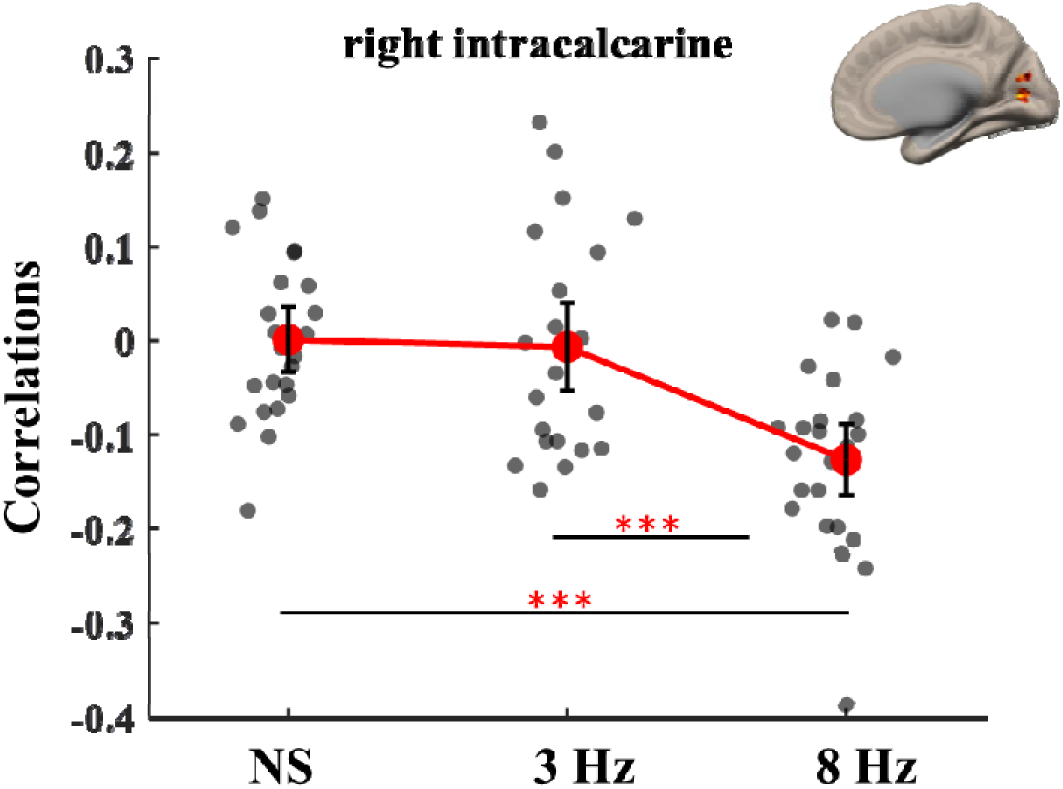
8 Hz modulation of P3-seeded functional connectivity with right visual cortex. Error bars represent the 95% confidence interval.

### Effect of tACS on hippocampal connectivity during retrieval

To test if tACS modulated hippocampal-cortical connectivity during memory retrieval, we applied the same hippocampal seed-based contrasts used for encoding (3 Hz contrast *c*=[0.5, −1, 0.5] and 8 Hz contrast *c*=[0.5, 0.5, −1]) to the temporal distance judgment task. Neither contrast yielded any voxel clusters surviving the significance threshold. These results indicate that, unlike during encoding, tACS did not significantly modulate hippocampal-cortical connectivity during retrieval, suggesting that the effects of slow theta stimulation on hippocampal network dynamics were specific to the encoding phase, when stimulation was actively applied.

## Discussion

In this study, we combined behavioral testing with concurrent tACS-fMRI to investigate how slow (3 Hz) versus fast (8 Hz) theta-frequency stimulation over the left parietal cortex influences hippocampal-cortical interactions during memory encoding. Behaviorally, we found that 3 Hz tACS led to shorter temporal distance judgments compared to 8 Hz stimulation, suggesting a frequency-specific impact on temporal memory organization. On the neural level, 3 Hz, but not 8 Hz, tACS reduced hippocampal functional connectivity with visual cortex during encoding. In addition, 3 Hz tACS reduced hippocampal activity but not visual cortex activity during encoding. Finally, we found no tACS-related connectivity effects during retrieval. These findings provide converging evidence that slow theta stimulation can modulate memory-relevant network dynamics and affect the encoding of temporal contextual information.

The observed hippocampal-cortical connectivity changes after 3 Hz stimulation included visual areas that have been implicated in visual scene processing and contextual representation (Altmann et al., 2004; Preston et al., 2013; Rehman & Al Khalili, 2025). Particularly, left lateral occipital cortex has also been shown to represent temporal contextual information about items encoded in contextual segments (Ezzyat & Davachi, 2014, 2021). Notably, in Ezzyat and Davachi (2014), visual items were shown on the right side of the screen and contextual information on the left side of the screen, similar to our task, which suggests that left visual cortex represented the temporal context of items, rather than identity of the contextual features. However, our findings suggest that 3 Hz tACS did not alter visual encoding per se, as we did not observe a change in visual cortical activity, but rather disrupted the integration of visual contextual information into hippocampal memory traces. In contrast, 8 Hz tACS did not significantly alter hippocampal-cortical connectivity, which is in line with previous findings of slow but not fast theta stimulation affecting memory performance (Aktürk et al., 2022; Wang et al., 2024; Wolinski et al., 2018). Further, it fits with our suggestion that 3 Hz tACS affected hippocampal functioning, as slow but not fast hippocampal theta has been found to be related to memory performance (Gray et al., 2026; Kota et al., 2020; Lega et al., 2012; Miller et al., 2018). We interpret this as an indirect effect of tACS via long-range projections at network-level connectivity beyond the stimulated site (Kaiser et al., 2025; Vieira et al., 2025), as the induced electric field at the scalp is too weak to directly affect neural activity in deep brain areas (Luo et al., 2025).

The observed decrease of temporal distance ratings could reflect strengthened associative or temporal binding during encoding. The perceived temporal compression of items encoded within a shared context may be due to the stronger associative encoding within an event than across boundaries (DuBrow & Davachi, 2013; Ezzyat & Davachi, 2014). Under this view, the compression observed under 3 Hz stimulation would indicate that slow theta tACS strengthened, rather than disrupted, the associative binding that ordinarily supports temporal proximity judgments. This interpretation would also explain why tACS enhanced associative memory false alarm rates without impacting hit rates, as the tendency for false alarms increases with increasing associative processing (Howe et al., 2009; Roediger & McDermott, 2000; van de Ven et al., 2018). This suggestion is also consistent with a previous report that slow theta tACS enhanced temporal memory accuracy (Wang et al., 2024). Another explanation is that slow theta tACS led to temporal compression because items appeared more similar than under fast theta stimulation. Temporal compression between items increases when the items are perceived to be more similar (Casasanto, 2008) and the pattern of hippocampal responses of individual items becomes more similar when the items are presented in the same context (Ezzyat & Davachi, 2014, 2021; Hsieh et al., 2014). Our observation of decreased hippocampal-visual cortical connectivity could indicate that fewer visual details were integrated into the memory traces, possibly increasing the similarity of the neural representations of those items. This interpretation could also explain the increased tACS-induced false alarm rates, which are also modulated by perceived item similarity. However, our experimental design of short and regularly timed item presentations prevented us from investigating how tACS modulated representational similarity in the brain. In addition, we note that the temporal distance rating used here is a measure of bias rather than accuracy, thereby limiting interpretation of mnemonic processing affected by tACS in our study. Nonetheless, if 3 Hz stimulation had impaired perceptual encoding or memory formation, ratings would be expected to regress toward the middle of the scale, reflecting greater uncertainty, or associative memory hit rates would be expected to decrease. As neither pattern of results was observed, we favor a mnemonic modulation interpretation, although we cannot rule out non-mnemonic effects of stimulation.

Our current finding differs from a previous report of increased resting-state hippocampal-parietal connectivity during theta tACS (Kaiser et al., 2025). This discrepancy may point to a dependency of tACS on brain state (Alagapan et al., 2016; Krause et al., 2022), where parietal connectivity with hippocampus may be more dominant during resting states while visual cortical connectivity may be more dominant during the encoding of visual item sequences (Ji & Wilson, 2007; Tambini et al., 2010). Moreover, the mnemonic component of our task may have further taxed hippocampal-visual cortical connectivity, as theta tACS during a visual task without a mnemonic component did not affect hippocampal-cortical connectivity (Kaiser et al., 2025). These considerations are in line with the more general suggestion that the effect of tACS on neural activity depends on brain state, as tACS temporally regroups endogenous responses rather than generates neural spiking *de novo* (Wischnewski et al., 2023). However, a more comprehensive description of the neural states underlying contextual encoding and event segmentation is required to obtain a better understanding of how tACS affects neural processing in hippocampus and cortex.

Finally, we found evidence for a partial dissociation between slow and fast theta frequencies delivered over the same parietal location modulating different neural pathways. That is, 3 Hz stimulation decreased hippocampal connectivity with left visual cortex while 8 Hz stimulation decreased parietal connectivity with right visual cortex, including early visual areas. The 8 Hz stimulation is spectrally close to the alpha band of 8-13 Hz, which has been amply associated with visual sensory and attentional processing (Bonnefond & Jensen, 2025; Clayton et al., 2018; Peylo et al., 2021; Zlatkova et al., 2026). The involvement of right (early) visual cortex during 8 Hz stimulation, as opposed to left (higher-order) visual cortex during 3 Hz stimulation, suggests that the two tACS frequencies impacted different functional components underlying the encoding task. That is, 3 Hz stimulation over parietal cortex may have affected hippocampal-cortical pathways because they utilize (slower) theta oscillatory activity to encode items in context. In contrast, 8 Hz stimulation over the same parietal site affected parietal-cortical pathways that are involved in sensory or attentional processing of the presented items. Anecdotally, it is striking that hippocampal connectivity includes higher-order visual cortex contralateral to the items presented with their contexts, while parietal connectivity includes early visual cortex contralateral to the presentation of the contexts themselves. Whether this lateralization indeed reflects item vs. context processing that affects segmentation and subsequent temporal memory formation remains to be investigated.

In conclusion, parietal slow theta (3 Hz) tACS modulated hippocampal-cortical connectivity during memory encoding and enhanced temporal compression. Our findings provide converging behavioral and neural evidence for a frequency-specific role of slow theta oscillations in shaping the temporal structure of episodic memory. Importantly, the observed effects were selective to encoding and were absent during retrieval, highlighting an online influence of tACS. These results advance our understanding of the neural dynamics underlying temporal memory and offer a foundation for developing network-targeted neuromodulation approaches in both healthy and clinical populations.

## Statements and declarations Author contributions

YW and VVDV designed the study, collected the data and analysed and visualized the data. YJ and LI contributed to data analysis and visualisation. FD and PDW contributed to study design. YW, VVDV and PDW secured project funding. YW wrote the first version of the manuscript. All authors contributed to manuscript writing and editing.

## Acknowledgments

We thank Tingting Zhu, Leia Bulgakova and Juanzhi Lu for their help in data collection.

## Funding

The study was partly funded by the Chinese Research Council (grant awarded to YW, PDW and VVDV) and MBIC-FPN scanning grants (awarded to YW and VVDV).

## Competing interests

The authors declare no competition of interests.

## Ethics approval

The ethics committee of the Faculty of Psychology and Neuroscience granted approval for the study, in line with the principles of the Declaration of Helsinki. All participants provided written informed consent prior to participation.

## Consent to publish

All authors provided consent to publish this work.

## Data availability

Behavioral data and compressed imaging data will be made available on a public server at acceptance of publication.

## Notes

### Competing Interest Statement

The authors have declared no competing interest.

